# Single cell multiomic analysis of the impact of Delta-9-tetrahydrocannabinol on HIV infected CD4 T cells

**DOI:** 10.1101/2025.06.02.657468

**Authors:** Manickam Ashokkumar, Renee Y Ge, Alicia Cooper-Volkheimer, David M Margolis, Quefeng Li, Yuchao Jiang, David M Murdoch, Edward P Browne

**Affiliations:** Department of Medicine, University of North Carolina at Chapel Hill, Chapel Hill, North Carolina, United States of America; UNC HIV Cure Center, University of North Carolina at Chapel Hill, Chapel Hill, North Carolina, United States of America; Department of Biostatistics, University of North Carolina at Chapel Hill, Chapel Hill, North Carolina, United States of America; Department of Medicine, Duke University, Durham, North Carolina, United States of America; Department of Statistics, Texas A&M University, College Station, Texas, United States of America; Department of Biology, Texas A&M University, College Station, Texas, United States of America; Department of Microbiology and Immunology, University of North Carolina at Chapel Hill, Chapel Hill, North Carolina, United States of America

**Author notes:** co-first or corresponding authors (Manickam Ashokkumar, Renee Y Ge, Yuchao Jiang, David M Murdoch, Edward P Browne).

## Abstract

Cannabis use is prevalent among individuals living with HIV in the United States, but the impact of cannabis exposure on the reservoir of latently infected cells that persists during antiretroviral therapy (ART) remains unclear. To address this gap, we analyzed the effect of Δ-9-tetrahydrocannabinol (THC) on primary CD4 T cells that were latently infected with HIV. We found that THC had no detectable effect on baseline or latency reversing agent (LRA) stimulated HIV expression, or on expression of an activation marker (CD38). However, using an integrated multiomic single-cell analysis of genome-wide chromatin accessibility and gene expression, we observed altered expression of several hundred genes in HIV infected CD4 T cells after THC exposure, including transcriptional downregulation of genes involved in protein translation and antiviral pathways, indicating that THC suppresses innate immune activation in infected cells. Additionally, chromatin accessibility analysis demonstrated upregulated chromatin binding activity for the transcriptional regulator CTCF, and reduced activity for members of the ETS transcription factor family in infected cells after THC exposure. These findings provide insights into the mechanisms by which cannabis use could influence the persistence of HIV within cellular reservoirs and the molecular phenotype of latently infected cells. Further elucidation of the underlying mechanisms involved in THC-mediated changes to HIV infected cells, will lead to an improved understanding of the impact of cannabis use on the HIV reservoir.

**Importance:** Cannabis use is common among individuals living with HIV, but the long-term effects of cannabis use on the HIV reservoir are not yet studied completely. We employed advanced single-cell technologies to reveal how cannabis components, specifically THC, influence HIV-infected immune cells and their pattern of gene expression. We found that, while THC doesn’t reactivate virus in latently infected cells, it alters the molecular characteristics of these infected immune cells. These findings are important because they underscore how cannabis could regulate persistent infection in people living with HIV. Understanding these cellular changes in response to THC could be helpful for successful treatment for people living with HIV.

## Introduction

Despite successful treatment with antiretroviral therapy (ART), people with HIV (PWH) retain a persistent viral reservoir that is the primary barrier to achieving a cure for HIV. This reservoir is comprised of a pool of latently infected cells that is distributed throughout various CD4 T cell subsets and tissues. In latently infected cells, viral gene expression is downregulated, but infected cells can still produce viral RNA and proteins which can potentially trigger an immune response and inflammation in PWH on ART (1). This persistently elevated immune activation can lead to a higher risk of non-AIDS comorbidities, including cardiovascular disease, cancer, kidney, liver, and neurologic diseases (2). Despite its importance, little is known about the latently infected reservoir, due the rare nature of the infected cells and the lack of a specific marker that allows purification and characterization of these cells.

Some evidence suggests that the immune system of PWH, as well as the size and nature of the reservoir, can be impacted by drugs of abuse. Cannabis (CB) use, in particular, is prevalent amongst people with HIV (PWH). We have recently reported that PWH on ART who use CB exhibit a trend towards a smaller intact HIV reservoir, as well as elevated frequencies of naïve T cells, and lower frequencies of activated and exhausted CD8 T cells. Consistent with this observation, another recent report indicated that CB use can lead to a reduced HIV reservoir burden in tissues in PWH (3). Other evidence from humans includes a recent study reporting that cannabis use was associated with lower frequencies of activated (HLA-DR^+^CD38^+^) CD4^+^ and CD8^+^ T cells, and significantly lower frequencies of TNF-α^+^ B cells in PWH on ART (2, 4). Animal studies have also pointed to an effect of CB on HIV infection and infection associated inflammation, including a recent report demonstrated that CB exposure leads to reduced inflammation in the gut of SIV infected animals (5). In a study conducted in a humanized mice model, researchers found that THC exposure increased the HIV viral load and decreased the number of IFN-γ secreting cells (6). By contrast, another study reported that chronic administration of THC to SIV infected rhesus macaques decreased viral load in cerebrospinal fluid (CSF) and plasma as well as mortality (7).

Cannabinoids are lipophilic molecules that have been shown to alter the functional activities of immune cells *in vitro* and *in vivo*. Δ-9-tetrahydrocannabinol (THC) is the most abundant and primary psychoactive cannabinoid component of cannabis. THC binds to and activates two endogenous cannabinoid receptors, CB1 and CB2. CB1 is largely present in brain and neurons, and CB2, by contrast, is expressed in immune cells, including CD4 T cells, the primary cell infected by HIV (8–10). Cannabinoid receptors regulate a complex signaling cascade that can affect CB-exposed cells on several levels. At the molecular level, activation of cannabinoid receptors leads to closure of Ca^2+^ channels and opening of K^+^ channels, inhibition of adenylyl cyclase activity and activation of a signaling cascade that leads to activation of the MAPK pathway (11, 12) including extracellular signal-regulated kinase 1/2 (ERK1/2), c-Jun N-terminal kinase (JNK), and p38, that are involved in the regulation of cell proliferation, cell cycle control and cell death. Cannabinoid receptors can also regulate expression of CREB and the pro-inflammatory transcription factor NF-κB (13).

CBs have complex effects on immune cell populations, and transcriptomic analyses of the effect of CBs on human immune cells have shown numerous alterations to transcriptional pathways in CD4 T cells (14). THC has been shown to suppress the activities of B and T lymphocytes, reduce the cytolytic activity of NK cells, inhibit the function and maturation of cytotoxic T lymphocytes (CTLs), and affect immune cell recruitment and chemotaxis to sites of infection (9). CBs are canonically thought of as immunosuppressive and reduce T cell proliferation and IL-2 expression after TCR-mediated activation, as well as TNF-α production and leukocyte migration across the blood brain barrier (9, 15). Thus, the collective data suggest that THC inhibits the functional activities of a variety of immune cells, an outcome that is consistent with these compounds altering host resistance to infectious agents (9).

Though the immunomodulatory properties of THC have been studied previously, its effect on cells that are latently infected with HIV has not yet been investigated. Since ∼50% of PWH use cannabis, it will be important to fully evaluate the impact of THC exposure on the HIV reservoir and its responsiveness to clinical latency reversal approaches. In this report, we use a primary CD4 T cell model of HIV latency and a single cell multiomic profiling approach to examine the impact of THC exposure on a population of latently infected cells. We find that, although THC does not have any apparent impact on HIV gene expression and responsiveness to latency reversing agents (LRAs) in this system, THC modulates the expression and activity of a set of genes and transcription factors in infected cells that may impact their behavior in PWH on ART.

## Methods

### Cell culture

Total CD4 T cells were isolated from peripheral blood mononuclear cells (PBMCs) from HIV seronegative donors using the EasySep Human CD4 T Cell Isolation Kit (STEMCELL Technologies, catalog # 17952) using negative selection, following the manufacturer’s protocol. Purity of CD4 T cells was determined by flow cytometry, which indicated high purity (>97%). Cells were seeded at 1 million cells/mL in RPMI media (Gibco), 10% fetal bovine serum (FBS), penicillin/streptomycin and incubated overnight at 37°C. A human embryonic kidney cell line that was used for transfection (HEK293T - ATCC; CRL11268) was maintained in DMEM complete medium (Gibco), supplemented with 10% FBS and penicillin/streptomycin.

### Primary CD4 T cell HIV latency model

Latently infected primary CD4 T cells were generated as described in our previous studies (16, 17). Briefly, CD4 T cells from seronegative donors were activated with anti-CD3/CD28 beads (Life Technologies) at 1M cells per mL for 3 days, then infected with a replication defective HIV reporter clone (NL4-3-△6-drEGFP-IRES-thy1.2) that had been pseudotyped with the Vesicular Stomatitis Virus G protein (VSV-G) (6, 23). Successful infection was confirmed by measuring GFP expression using flow cytometry followed by enrichment of GFP+ infected cells using a FACSAriaIII (Becton Dickson). Enriched infected cells were then kept in culture for 3 weeks while they returned to a resting state and viral gene expression was downregulated. The infected CD4 T cells and matched PBMCs were then mixed at a ratio of 1:3 to generate a mixed population of latently infected cells and PBMCs.

### Δ-9-tetrahydrocannabinol stimulation and LRA treatment

Latently infected primary CD4 cells were stimulated with Δ-9-tetrahydrocannabinol (THC) at concentrations ranging from 0μM to 10μM for 6 h or empty vehicle (methanol). At 6 h post THC exposure, the cells were harvested, washed and quantified for viral GFP expression, CD38 cell activation marker, and CD4 surface marker expression by flow cytometry. In addition to the dose response stimulation with THC, we also quantified reactivation and cell activation by measuring GFP and CD38 in different CD4 T cell subsets with THC exposure followed by LRA stimulation by one of three different LRAs with different mechanisms of action - AZD5582 (non-canonical NF-κB agonist), prostratin (PKC agonist), or vorinostat (HDAC inhibitor) or control vehicle (DMSO).

### THC stimulation for single cell ATAC and Gene expression

HIV infected primary CD4 T cells pooled with PBMCs of the same donor were stimulated with Δ-9-tetrahydrocannabinol (THC) at a concentration of 500nM or control vehicle (methanol 0.015%). Cells were then combined with autologous PBMCs that had been thawed and rested for 24h at 37C at a ratio of 1:3 (CD4 T cells : PBMCs). Each experiment was carried out with three biological replicates (separate donors). At 6 h post THC exposure, the cells were harvested for integrated single cell multi-omic ATAC and gene expression library preparation.

### Nuclei isolation

Nuclei were isolated following the 10xGenomics protocol (CG000365 • Rev C) as described in our previous study (17). Cells were harvested, washed once with ice-cold phosphate buffered saline (PBS)-0.5% BSA containing 0.2 U/μl RiboLock RNase inhibitor (Thermo Fisher, cat. no. EO0382), counted and viability determined by Trypan blue exclusion (Thermo Fisher, cat. no.15250061). Dead cells were removed using a dead cell removal kit (Miltenyi Biotec, cat. no. 130-090-101. For each sample, 1×10^6^ cells were collected by centrifugation at 500 g, 4°C, 5 min. The cells were then resuspended in 100 μl of ice-cold lysis buffer (10 mM Tris-HCl, pH 7.4, 10 mM sodium chloride, 3mM magnesium chloride, 0.01% Tween-20 (Sigma, cat. no. 655205-250ML), 0.01% nonidet P40 substitute (Sigma, cat. no. I8896), 0.01% Digitonin (Promega, cat. no. G9441), 1 U/μl RiboLock RNase inhibitor, 1% BSA and 1mM DTT) and incubated on ice for 3 min. The released nuclei were then washed with 1 ml ice-cold wash buffer (10 mM Tris-HCl, pH 7.4, 10 mM sodium chloride, 3 mM magnesium chloride, 0.01% Tween-20, 1% BSA, 1mM DTT and 1 U/μl RiboLock RNase inhibitor) and nuclei were re-suspended in diluted nuclei buffer (1X nuclei buffer, 1mM DTT, 1 U/μl RiboLock RNase inhibitor).

### Construction of integrated single cell ATACseq and gene expression libraries and sequencing

Nuclei samples were used to generate single cell ATAC-seq and RNA-seq libraries using a Chromium Single Cell Controller (10xGenomics, Pleasanton, CA) and a Single Cell multiome ATAC and gene expression kit (CG000338 • Rev E). Briefly, diluted nuclei suspensions with a target recovery of 10,000 nuclei were subjected to Tn5 transposition in bulk, followed by barcoding using GEM (Gel Beads-in-emulsion) beads. Silane magnetic beads were used to purify the barcoded products from the post GEM-RT reaction mixture. Barcoded ATAC fragments from the transposed DNA and barcoded, full-length cDNA from poly-adenylated mRNA were then pre-amplified by polymerase chain reaction (PCR) to facilitate library construction. The pre-amplified product was then used as input for both ATAC and gene expression (GEX) library construction. P5 and P7 indices were added to the pre-amplified transposed DNA for ATAC-seq library.

cDNA amplification, enzymatic fragmentation followed by end repair, A-tailing, adaptor ligation, and PCR were performed to incorporate P5, P7, i7 and i5 sample indices, and TruSeq Read 2 (read 2 primer sequence) for gene expression libraries. The libraries were quantified using an Agilent Tapestation 4200 and the Qubit dsDNA High Sensitivity Assay Kit (Invitrogen, #Q33230). Pooled samples for the ATAC and RNA libraries were sequenced using paired-end, single-index (ATAC-seq) and dual index (RNA-seq) sequencing on a NextSeq 2000 instrument (Illumina). For GEX libraries, the read format was: Read 1 – 28 cycles; Read 2 – 90 cycles; i7 – 10 cycles; i5 – 10 cycles. For ATAC libraries the read format was: Read 1 – 50 cycles; Read 2 – 49 cycles; i7 – 8 cycles, i5 – 16 cycles. Paired end reads of pooled libraries were demultiplexed prior to downstream analysis.

### scRNA-seq and scATAC-seq data processing

Data from the three separate donors were first processed separately. Cells with high percentages of mitochondrial reads, extreme RNA read counts, and/or extreme ATAC fragment counts were removed. Genes and peaks not expressed/accessible in at least 0.5% of the cells were removed with the exception of the HIV genes. Doublet cells, identified by scDblFinder (18), were excluded from subsequent analysis. For read count normalization, we used sctransform by Seurat (19) for scRNA-seq and term frequency-inverse document frequency (TF-IDF) by Signac (20) for scATAC-seq. This was followed by principal component analysis (PCA) and latent semantic indexing (LSI) for dimension reduction, respectively. Cell-type labels were transferred from an existing curated and annotated PBMC reference dataset (21); cells with maximum prediction scores less than 0.8 for cell-type label transferring were removed. For modality-specific processing, we built the unweighted K-nearest neighbor (KNN) and the weighted shared nearest neighbor (SNN) graphs, followed by UMAP visualization and clustering identification (22).

To account for batch effects between the three biological replicates in the scRNA-seq data, we used anchor-based CCA Integration (23) to produce a joint dimension reduction, which was further used for UMAP visualization and clustering. Based on the integrated RNA UMAP, we observed CD4 T cells in two main subclusters: one exhibiting HIV Expression (HIV+ CD4 T) and one without HIV Expression (HIV-CD4 T). For analyses regarding the impact of THC on HIV infected cells, only cells from the HIV+ CD4 HIV cluster were considered. Clusters containing less than 50 cells were removed. Within the scATAC-seq data, we integrate the LSI embeddings across the donors, followed by UMAP visualization and clustering. We obtained the position frequency matrices and annotated 633 TF-binding motifs from the JASPAR 2020 database (24); we further applied chromVAR (25) to derive, for each TF, its motif deviation score, which measures the deviation in chromatin accessibility across the set of peaks containing the corresponding motif, compared with a set of background peaks.

For each cell-type cluster, we performed a nonparametric Wilcoxon rank sum test to identify differentially expressed genes (DEGs) and differentially accessible motifs between the THC- and THC+ conditions. We used the normalized gene expression levels by sctransform and the motif deviation scores by chromVAR as input, and adopted false discovery rate (FDR) for multiple testing correction. For significant DEGs, we carried out a gene ontology (GO) enrichment analysis using Enrichr (26).

## Results

### Impact of THC on CD4 T cell surface marker phenotype and HIV expression

Although we and others have previously shown that cannabis (CB) use is associated with an altered abundance of T cell subsets and expression of activation markers in people with HIV (PWH) on antiretroviral therapy (ART), it is unknown if this reflects a direct impact of CB on the immune cells or on expression of the HIV reservoir. Therefore, we examined whether THC, the primary cannabinoid present in CB, affects the phenotype of HIV infected cells or expression of HIV using a primary CD4 T cell latency model that we have previously established. We added THC over a range of concentrations (0-10μM) to *ex vivo* generated latently infected primary CD4+ T cells (**Figure 1A**). At 6h and 24h post stimulation with THC, flow cytometry was performed to determine the abundance of CD4 T cell subsets based on key surface markers (CCR7, CD45RA, and CD38), and HIV expression (GFP). We first used CCR7 and CD45RA staining to assign cells as naïve (Tn - CD45RA+/CCR7+), central memory (Tcm - CD45RA-/CCR7+), effector memory (Tem - CD45RA-/CCR7-) or terminally differentiated effector cells (Teff - CD45RA+/CCR7-). We observed that, overall, *in vitro* THC exposure of latently infected primary CD4+ T cells did not alter the frequency of cell subsets significantly at either timepoint (**Figure 1B, and Figure S1A**). We then examined the impact of THC on expression of the surface marker CD38 in HIV infected CD4 T cells. CD38 is frequently used as a marker for activated CD4 T cells, but is also constitutively expressed in naïve and central memory CD4 T cells, where it regulates several aspects of CD4 T cell function, including regulating cellular NAD+ levels (27). We observed no change in surface expression of CD38 for any of the subsets at any THC concentration (**Figure 1C**). These data indicate that THC exposure does not have a detectable impact on the expression of CD4 T cell subset surface markers or on expression of CD38.

**Figure 1.**
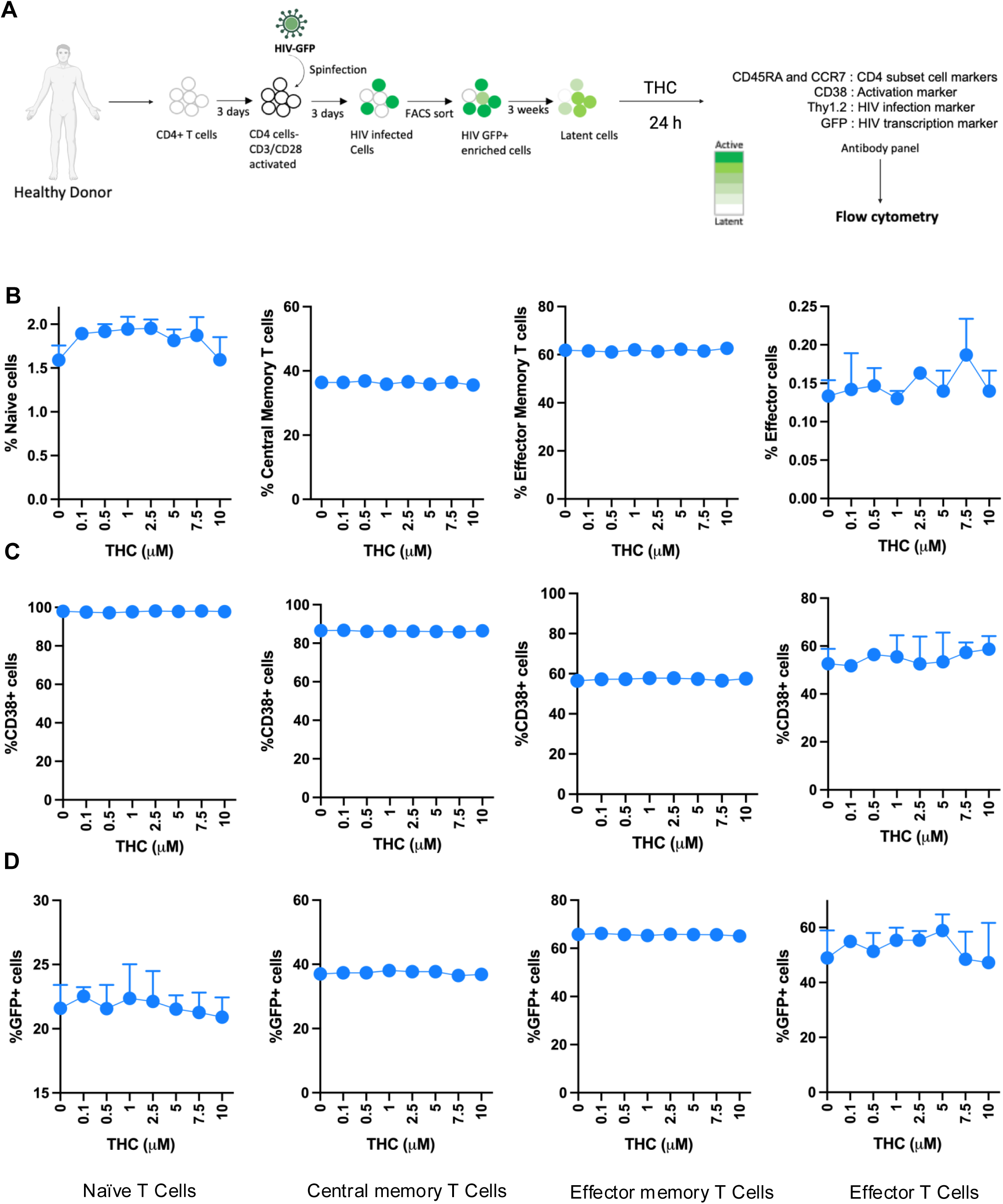
Impact of THC on cell surface marker phenotype and HIV expression in latently infected primary CD4 cells. Latently infected CD4 cells were exposed to THC at different concentration for 24 h. At 24 h post exposure, we quantified the distribution of different CD4 T cell subsets, expression of activation marker, CD38, and HIV (GFP) expression. Cell subsets were assigned by expression of the phenotype surface markers, CD45RA and CCR7 – Tn: CD45RA+/CCR7+, Tcm: CD45RA-/CCR7+, Tem: CD45RA-/CCR7-, Teff: CD45RA+/CCR7-. (**A**) Schematic of latently infected CD4+ T cell generation, pooling and THC stimulation followed by flowcytometry. (**B**) Abundance of CD4 T cell subsets - naïve (Tn), central memory (Tcm), effector memory (Tem) and effector (Teff) subsets at 24 h post THC administration/exposure. (**C**) Frequency of expression for the immune surface marker CD38 within CD4 T cells subsets (Tn, Tcm, Tem, and Teff cells). (**D**) Expression of eGFP (HIV) within CD4 T cells subsets (Tn, Tcm, Tem, and Teff cells). None of the samples achieved statistical significance relative to unexposed control cells.

We then examined the impact of THC on viral gene expression (GFP) in infected cells (**Figure 1D**). In our latency model system, CD4 T cells are first activated, then infected with a GFP-expressing strain of HIV (HIV-GFP). Actively infected cells are then flow sorted to obtain a pure infected population (GFP+). These cells are then cultured for three weeks, during which time viral gene expression is progressively downregulated and a latently infected (GFP-) population emerges, although residual GFP expression remains for a subset of the infected cells. When we examined the level of baseline viral gene expression across the different T cell subsets, we observed that Tn and Tcm exhibited a lower level of baseline HIV expression than Tem and Teff cells (21.6% and 37% v 65.8% and 48.9%). Notably, expression of HIV was not affected by the presence of THC at any concentration in any of the CD4 T cell subsets (**Figure 1D**). We also examined the impact of THC on the subset markers, CD38 expression and viral GFP expression at an earlier timepoint (6h) and found no impact of THC on any of these parameters at any THC concentration (**Figure S1**). Overall, these data indicate that, in this *ex vivo* latency model system, THC does not have a major impact on the maturation status, activation status or baseline HIV expression in infected CD4 T cells.

### Reactivation of HIV from latency by latency reversing agents is not impacted by THC

A major approach in eliminating latently infected cells from PWH on ART involves the pharmacological reactivation of HIV expression with small molecules referred to as latency reversing agents (LRAs). To assess the impact of THC exposure on reactivation of latently infected CD4 T cells by LRAs, we stimulated latently infected CD4 T cells with THC (0, 0.5, 1 and 10μM) for 24 h followed by a 24 h exposure to one of three different LRAs with different mechanisms of action - AZD5582 (non-canonical NF-κB agonist), prostratin (PKC agonist), or vorinostat (HDAC inhibitor) or control vehicle (DMSO). The cells were then analyzed by flow cytometry to measure expression of CD4 T cell subset surface markers (CCR7, CD45RA, CD38), and viral gene expression (GFP). To facilitate analysis and comparison across cell types, datasets were converted into fold change value normalized to the control vehicle condition. First, we compared the relative frequencies of the CD4 T cell subsets (Tn, Tcm, Tem and Teff) following LRA stimulation with THC exposure. Interestingly, we noticed that some of the LRAs affected the expression of the CD4 T cell subset markers (**Figure 2A**). Specifically, we observed that AZD5582 increased the frequency of cells with a Tcm phenotype and decreased the proportion of cells with a Tn or Teff phenotype, while prostratin exposure decreased Tcm cells and increased Teff cells. By contrast, vorinostat had no impact on the expression of subset surface markers. As before, we found that THC exposure had no effect on the frequencies of the different T cell subsets in any of the LRA stimulated conditions. Next, we sought to examine the impact of combined LRA and THC exposure on expression of CD38 in latently infected CD4 T cell subsets (**Figure 2B**). LRA exposure did not strongly affect CD38 expression for most CD4 T cell subsets, although we observed a modest decrease for CD38 expression in Tcm cells exposed to AZD5582, and a modest increase in CD38 for Tem cells exposed to prostratin. When we compared expression of CD38 across the different THC concentrations and LRA conditions, we observed no impact of THC at any concentration in any cell subset. Thus, THC does not affect CD4 T cell subset marker or CD38 expression in the presence or absence of LRAs.

**Figure 2.**
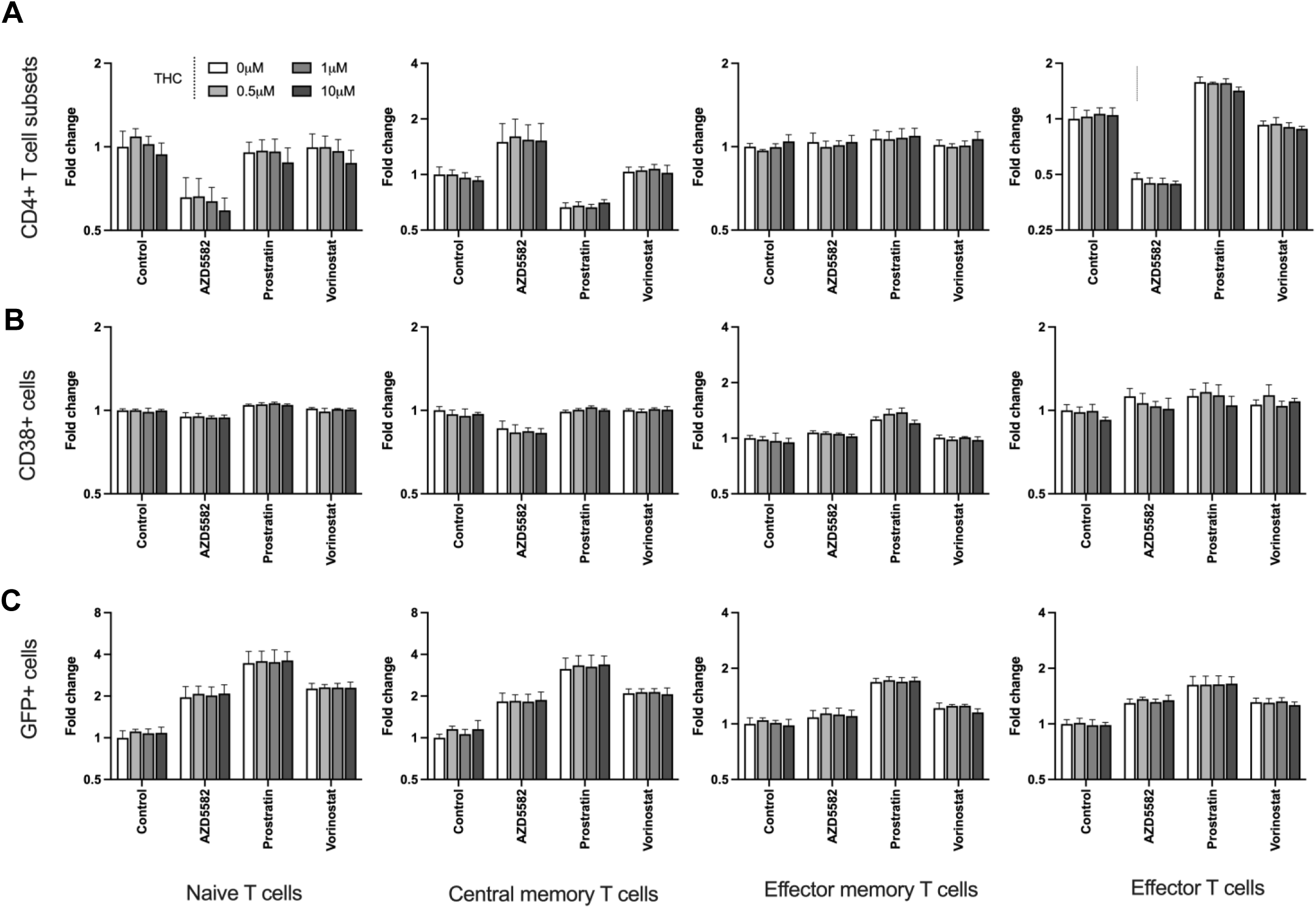
Impact of THC on viral reactivation by latency reversing agents. To assess the impact of THC exposure on reactivation of latently infected CD4 T cells by latency reversing agents (LRAs), we stimulated latently infected CD4 T cells with THC (0, 0.5, 1 and 10μM) for 24 h followed by a 24 h exposure to one of the three different LRAs with different mechanisms of action - AZD5582 (non-canonical NF-κB agonist), prostratin (PKC agonist), and vorinostat (HDAC inhibitor) or control vehicle (DMSO). (**A**) Abundance (fold change) of CD4 T cell subsets (Tn, Tcm, Tem, Teff) exposed to THC followed by LRAs stimulation. (**B**) Expression level (fold change) of CD38 is shown for latently infected CD4 T cell subsets (Tn, Tcm, Tem, and Teff cells) exposed to THC followed by LRAs stimulation. (**C**) Expression level (fold change) of GFP is shown for latently infected CD4 T cell subsets (Tn, Tcm, Tem, and Teff cells) exposed to THC followed by LRAs stimulation. None of the bars/sample reached statistical significance compared to control vehicle.

Next, we examined HIV reactivation by LRAs in different CD4+ T cell subsets in the presence of THC. After stimulation with the LRAs, we observed that Tn and Tcm exhibited the highest fold change increase in GFP+ cells over baseline, while LRA responses in Tem and Teff were lower, likely due to their lower level of restriction to HIV expression (**Figure 2C**). Overall, prostratin was consistently the most potent LRA in all of the CD4 T cell subsets. We observed that THC exposure had no impact on latency reversal for any of the LRAs in any of the CD4 T cell subsets, suggesting that HIV expression and reactivation is robust to THC exposure.

### Integrated Single cell RNA- and ATAC-seq profiling of THC stimulated HIV infected cells

Although our data indicated that THC exposure has little impact on latently infected cells in terms of surface marker expression or HIV expression at the protein level, we speculated that THC could have an impact on aspects of the host cells not detectable by these assays. Since most immune cells, including CD4 T cells, express cannabinoid receptors, we hypothesized that THC exposure triggers definable changes to the transcriptome of infected cells that could impact their biological properties. To investigate this hypothesis, we stimulated a mixture of latently infected CD4 T cells and autologous PBMCs (at a ratio of 1:3) from three independent donors with 500nM THC. To capture immediate early transcriptional responses to stimulation, we profiled these cells at 6 hr post exposure for changes in accessible chromatin and gene expression using a combined single cell RNAseq/ATACseq approach (**Figure 3A**). After quality control and batch effect removal, integrated scRNA-seq/scATAC-seq yielded a total of 18,407 cells for pre- and 20,956 cells for post-THC exposure with detectable expression of ∼15,000 genes and ∼110,700 accessible chromatin peaks detected across the cell population.

**Figure 3.**
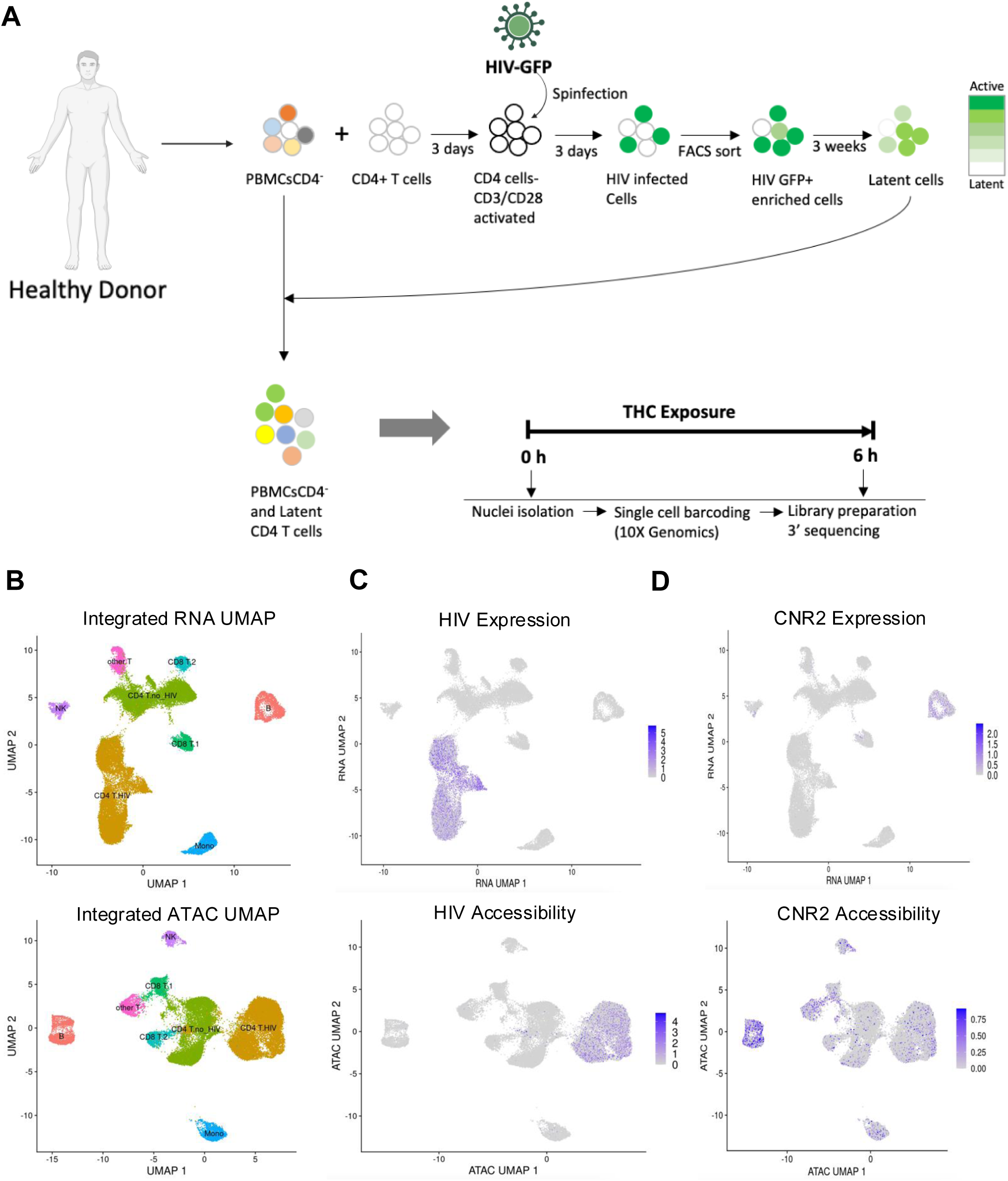
Single cell multiomic analysis of latently infected CD4+ T cells exposed to THC. (**A**) Schematic of experimental design. (**B**) Uniform Manifold Approximation and Projection (UMAP) dimension reduction of scRNAseq (top) and scATACseq (bottom) with cells labeled by immune cell type. (**C**) UMAP plot of scRNAseq (top) and scATACseq (bottom) data with cells containing HIV mapping RNA or ATAC reads labeled purple. (**D**) Same as C but with cells containing RNA or ATAC reads from the CNR2 gene labeled purple. Data show represents aggregated data from three independent donors.

Dimension reduction using Uniform Manifold Approximation and Projection (UMAP) of both the scRNAseq and scATACseq data showed several distinct clusters of immune cells (**Figure 3B**). To annotate the cell types of the clusters, we used a generalized linear model (GLM)-based cell mapping approach with cell-type “marker” genes curated from the available literature. Briefly, we selected a reference gene panel based on known cell-type-specific gene profiles, then used GLM to test the association of gene expression in each cell with the known marker genes (**Figure 3B**). Through the expression of marker genes, each cluster was assigned to a cell type based on the highest percentage of significant cells. This approach deconvoluted the 39,363 cells into specific cell subtypes: B cells, NK cells, Monocytes, CD4 T cells and CD8 T cells. CD4 and were divided into two separate clusters each, with one cluster likely reflecting the cells that were activated, infected and returned to rest (“CD4 T HIV”) and other being the cells that were added back to the culture after three weeks as part of the PBMC population (“CD4 T no HIV”). CD8 T cells were also present as two major clusters (“CD8T1” and “CD8 T2”), possibly due to CD8 T cells that contaminated the CD4 T cell population that was activated and infected. Another subpopulation of T cells remained unclassified (“other T cells”).

Next, we examined the distribution of viral RNA and ATAC reads across the clusters. As expected, HIV-mapping reads were found largely within only one of the two CD4 T cell cluster sets (CD4 T HIV) for both viral RNA and ATAC reads, allowing us to identify the CD4 T cells derived from the HIV infected culture (**Figure 3C**). These data demonstrate that we can identify cells infected with HIV from the presence of reads that map to the viral genome. We also examined the expression and accessibility pattern for the CNR2 gene which encodes the CB2 THC receptor. Interestingly, at the RNA levels CNR2 expression was highest in B cells, while the CNR2 gene was accessible across most immune cell types (**Figure 3D**).

### Impact of THC on viral transcription and proviral accessibility

We next examined the impact of THC exposure on the HIV proviruses in infected cells at the level of viral transcription and proviral accessibility. During transcriptional activation, pioneer transcription factors recruit chromatin remodeling complexes to regulatory regions, resulting in increased accessibility at these regions, and these changes in chromatin accessibility are known to play a crucial role in transcription (28). The combined scRNAseq/scATACseq approach thus allows us to examine the relationship between changes in accessibility and expression for individual genes. For each cell, we quantified the number of viral RNA (vRNA) reads and viral ATAC (vATAC) reads and we examined the correlation between vRNA and vATAC reads across the cell population (**Figure 4A**). Consistent with our previous findings (17), we observed a significant positive correlation between the frequency of vRNA and vATAC reads within the cells, and this relationship was also true for both THC-stimulated and unstimulated cells alone (**Figure 4A**). We examined the pattern of vRNA and vATAC reads across the CD4 T HIV UMAP plot, we observed that THC exposure did not noticeably change the pattern of reads distribution for either vRNA or vATAC reads (**Figure S2)**. We then examined the proportion of CD4 T cells with vRNA and vATAC reads as well as the average abundance of vRNA (**Figure 4B**) and vATAC (**Figure 4C**) reads across the population. Interestingly, we observed that the scRNAseq and scATAC-seq reads mapping to the HIV genome showed a small but statistically significant reduction in vRNA expression and chromatin accessibility for the HIV infected CD4 T cell population exposed to THC. Thus, THC may have a small inhibitory effect on HIV transcription in latently infected cells. Nevertheless, we conclude that, overall, THC has only a minor impact on HIV expression and accessibility in latently infected cells, consistent with our observations by flow cytometry.

**Figure 4.**
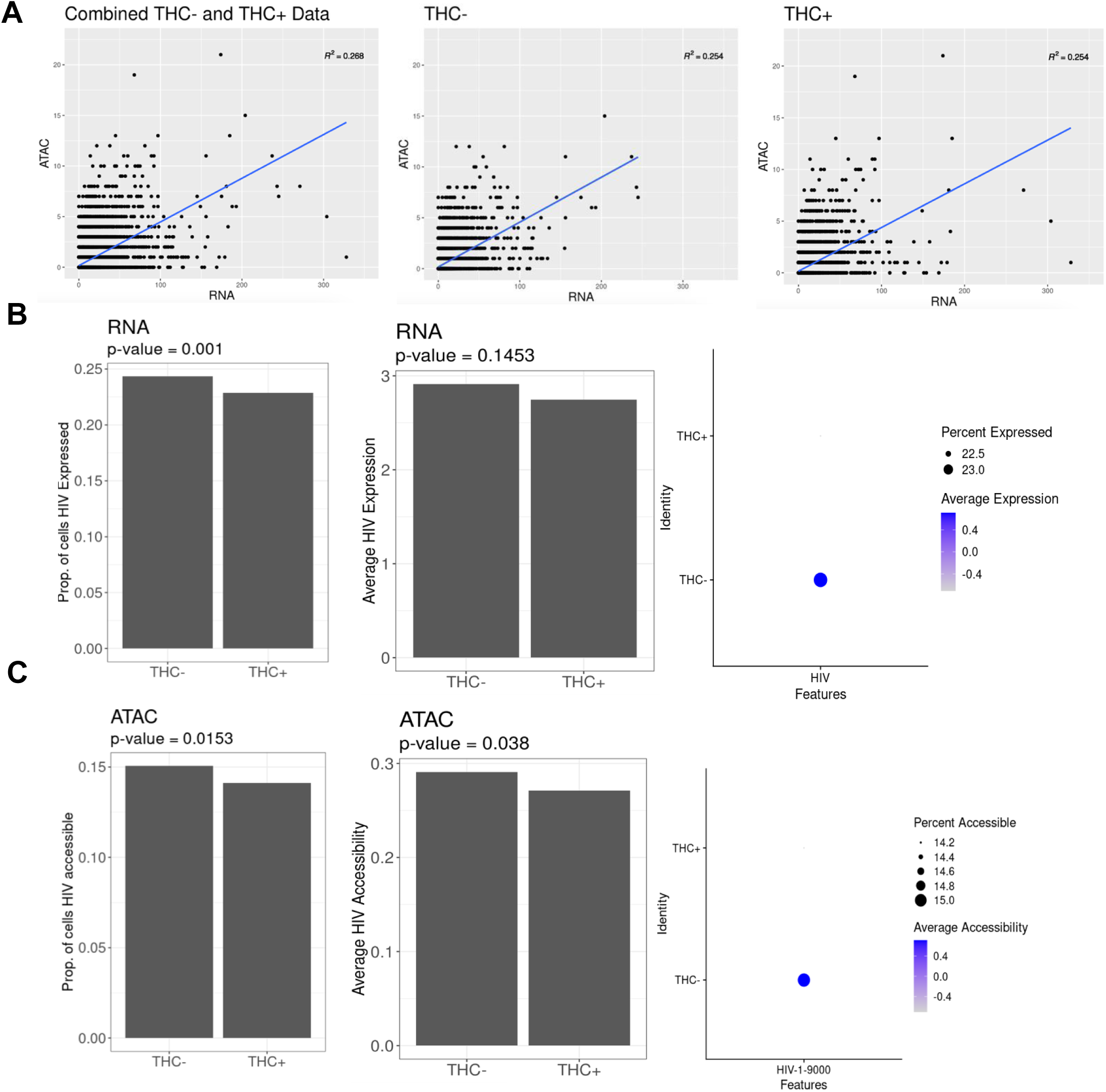
HIV vRNA expression and chromatin accessibility is mildly affected by THC exposure. **(A)** Correlation scatter plot showing the square root (sqrt) proportion of scRNAseq reads mapping to HIV (X-axis) vs. the sqrt proportion of scATACseq reads mapping to HIV (Y-axis) across the cell population for the integrated data (Left panel), without THC exposure (Middle panel), and with THC exposure (Right panel) for 6 h. **(B)** Transformed HIV expression data (proportion of cells accessible to HIV, Left panel; Average HIV accessibility, Middle panel; and Dot plot for percentage and average accessibility, Right panel) for cells without and with exposure to THC. **(C)** Transformed HIV ATAC accessibility data (proportion of cells accessible to HIV, Left panel; Average HIV accessibility, Middle panel; and Dot plot for percentage and average accessibility, Right panel) for cells without and with exposure to THC. A permutation non-parametric statistical method were used to assess the significance.

### THC exposure modulates infected host cell transcription

Although we observed only a minor impact of THC exposure on HIV expression and accessibility, we hypothesized that THC could impact the transcriptomic or epigenomic phenotype of infected cells as well as uninfected cells within the culture. In addition to allowing us to identify genes whose expression is affected by a given stimulus, the combined scRNAseq/scATACseq approach allows us to quantify the activity of cellular transcription factors (TFs) within each cell based on the enrichment of TF binding sites within accessible chromatin regions.

To maximize our ability to identify THC-induced changes in gene expression and TF activity, we considered DEGs within specific subsets of cells (B cells, NK cells, monocytes, CD8 T cells, CD4 T cells). For CD4 T cells we also separately considered cells from the HIV infected cluster and the uninfected (PBMC derived) cluster. Across the different immune cell types, we observed considerable variation in the magnitude of the transcriptomic response to THC (**Figure 5A, Table S1**). In most cell subsets, we observed a small number of DEGs, with 15 DEGs in B cells (11 up, four down), two DEGs in NK cells, two DEGs in monocytes, seven DEGs in CD8 cluster 1 T cells and 37 in CD8 cluster 2 T cells. By contrast, CD4 T cells exhibited a more pronounced THC response, with 271 DEGs in the uninfected CD4 T cell cluster and 515 DEGs in the HIV infected CD4 T cell cluster. Notably, the changes in gene expression for both the CD4 T cell clusters were highly polarized, with 35 upregulated genes and 236 downregulated genes in uninfected CD4 T cells, and 22 upregulated and 493 downregulated genes in the infected cells (**Figure 5A**). These data suggest that THC exposure has an overall suppressive effect on gene expression in CD4 T cells.

**Figure 5.**
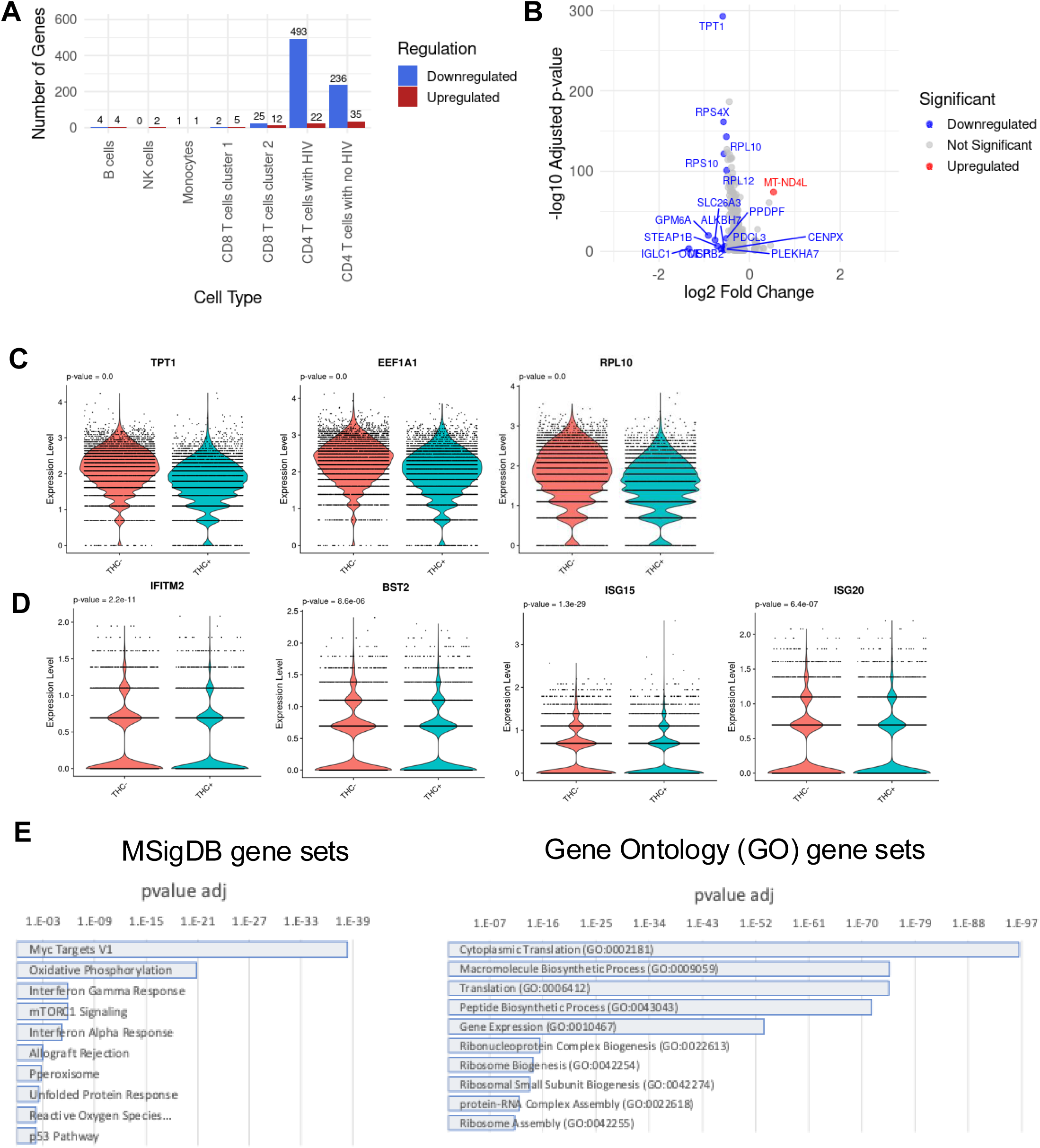
THC affects gene expression in HIV infected CD4 T cells. (A) Bar plot showing the number of differentially expressed genes (DEGs) across different immune cell types. **(B)** Volcano plot of overall differentially expressed genes (DEGs) between HIV-1 infected CD4 T cells exposed to THC vs non-exposure. The significance in DEGs were set with the log2 fold change threshold <-0.5 and the adjusted p-value threshold <-0.05. **(C)** Violin plot showing downregulation of selected genes that contribute to protein translation, including genes encoding ribosomal subunits (TPT1, EEF1A1, and RPL10) in response to THC exposure. **(D)** Violin plot showing expression of selected interferon sensitive genes (IFITM2, ISG15, ISG20 and BST2) in THC-stimulated HIV infected cells. P values from Wilcoxon rank sum tests comparing control and THC exposure. **(E)** MSigDB gene set enrichment (Left panel) and Gene ontology analysis of downregulated DEGs for HIV infected cells after THC exposure.

Several noteworthy genes were observed in the set of differentially expressed genes (**Table S1**). In B cells and in uninfected CD4 T cells, we observed downregulation of the pro-inflammatory cytokine IL-1β, consistent with previous reports that CB signaling inhibits inflammasome activation (29). In THC stimulated B cells, CD8 T cells and uninfected CD4 T cells we observed strong downregulation of the ion transporter SLC26A3 that regulated intestinal permeability. Within both infected and uninfected CD4 T cells, the set of downregulated DEGs contained numerous genes that contribute to protein translation, including genes encoding ribosomal subunits, suggesting a negative impact of THC on overall cell metabolism and protein synthesis in infected cells (**Figure 5C**). Interestingly, we also observed that several known interferon stimulating genes (ISGs) were downregulated in THC-stimulated HIV infected cells including IFITM2, ISG15, ISG20 and BST2 (**Figure 5D**). Notably, BST2 has been identified as a key HIV restriction factor that inhibits HIV particle release and is counteracted by the viral protein, Vpu. These data suggest that THC exposure could contribute to suppressing the expression of innate antiviral pathways in infected cells.

We then examined enrichment of specific biological pathways in the sets of DEGs for uninfected and infected CD4 T cells. Specifically, we examined the DEGs for enrichment with curated sets of pathway-associated genes using ENRICHR (26). For both infected and uninfected CD4 T cells, no specific pathways were found to be enriched in the set of THC-upregulated genes, likely due to the small numbers of upregulated DEGs. By contrast, in the set of genes that were downregulated in uninfected CD4 T cells, we observed significant enrichment for four genes sets within MySIGDB - Myc target genes, TNF-alpha Signaling via NF-κB, p53 Pathway, and Allograft Rejection (**<u>Table S2</u>**). In infected cells, 19 pathways were enriched, including Myc Targets, mTORC1 signaling and oxidative phosphorylation (**Figure 5E left panel, Table S3**). For the GO Biological Process sets, Cytoplasmic translation (GO:0002181) and Macromolecule Biosynthetic Process (GO:0009059) were the top two enriched gene sets in the downregulated DEGs for both infected and uninfected cells, consistent with the hypothesis that THC has a negative effect on protein synthesis and metabolism for both infected and uninfected cells (**Figure 5E right panel, Table S4, Table S5**).

### THC exposure affects the activity of specific TFs in HIV infected CD4 T cells

We next examined the impact of THC exposure on the activity of transcription factors within the different immune cell populations. During transcriptional activation, pioneer transcription factors bind to DNA in a sequence-specific manner and recruit chromatin remodeling complexes that promote increased accessibility within key gene controlling regions (28). Since scATACseq data provides genome-wide accessibility information from each cell, we can thus estimate the activity of cellular transcription factors within each cell from these data. For each cell in the population, a TF activity score was calculated for a set of 600 cellular TFs with known binding site sequences, based on the enrichment of these sequences within the open chromatin in that cell. Using this TF activity matrix we then identified differentially active transcription factors (DATFs) that exhibited altered activity after THC stimulation (**Table S6**).

Similar to our observations with differentially expressed genes (DEGs), we observed that the impact of THC was highly variable depending on the immune cell type (**Figure 6A, Table S6**). Surprisingly, no DATFs were identified in B cells, despite high expression of CNR2 in these cells. Six THC-responsive DATFs were identified in monocytes – one that was upregulated (FIGLA) and five that were downregulated (GABPA, ELF1, ELF3, EHF, and IKZF1). Notably, all of the downregulated DATFs except IKZF1 are members of the ETS TF family. Within NK cells, a single DATF was identified, the transcriptional regulator CTCF, whose activity was increased in NK cells in response to THC exposure. In CD8 T cells, we observed 7 DATFs, all which were downregulated and, notably, all of these DATFs were members of ETS1 family.

**Figure 6.**
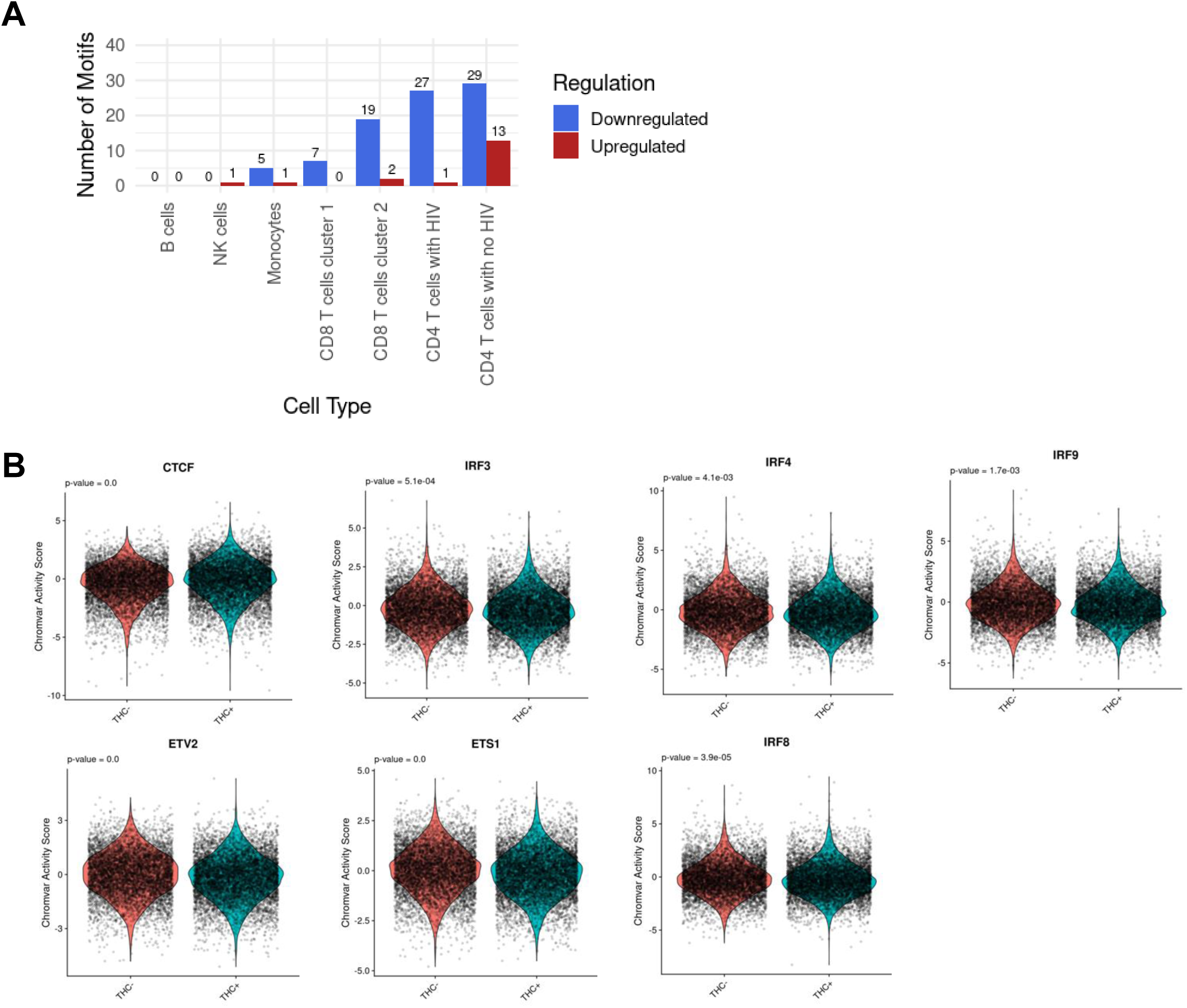
THC impacts transcription factor activity in HIV infected CD4 T cells. (A) Bar plot showing the number of differentially active transcription factors (DATFs) across the different immune cell types after 6 h THC exposure. **(B)** Violin plots showing activity scores for selected TFs (CTCF, IRF3, IRF4, IRF9, ETV2, ETS1, IRF8) in response to THC exposure. P values from Wilcoxon rank sum tests comparing THC treated cells to control treated cells.

Consistent with our DEG analysis, we observed that the strongest impact of THC was within CD4 T cells. Within uninfected CD4 T cells, we observed 42 DATFs - 13 upregulated and 29 downregulated. Similar to our observations with monocytes, the downregulated DATFs were largely composed of ETS TFs, but also included IKZF1. Within the set of upregulated DATFs, CTCF was the most highly upregulated, although two TFs that have been shown to regulate HIV expression, ATF4 and YY1, were also upregulated. Within HIV infected CD4 T cells, we observed 28 DATFs, only one of which (CTCF) was upregulated. Similar to what we had observed for uninfected CD4 T cells, the majority of the 27 downregulated TFs were ETS family members (ETV2, ETS1, ERG, FLI1, ELF4, ELK3, ERF, FEV, ETV3, ETS2, ELF5, EHF, ETV5, ELF2, ETV6, ELK4, ELF3, GABPA, ELF1, ELK1, ETV1), while IKZF1 was also downregulated. Interestingly, however, we also observed that there were three interferon regulated TFs (IRF3, IRF8 and IRF9) that were downregulated in infected CD4 T cells (**Figure 6B, Table S6**), but not in uninfected CD4 T cells. This finding is consistent with our observation that ISGs were transcriptionally downregulated in HIV infected CD4 T cells. Overall, these data indicate that THC exposure affects the activity of several TFs in HIV infected CD4 T cells. In particular, we consistently observed that CTCF activity was upregulated while ETS TFs and IRFs were downregulated in THC exposed HIV infected CD4 T cells. These altered patterns of TF activity likely explain the altered gene expression in these cells after THC exposure.

## Discussion

Cannabis (CB) use is common in people with HIV (PWH) and CB exposure may exert potential therapeutic benefits to PWH due to its anti-inflammatory properties (30, 31). It was previously reported that cannabis use is associated with a lower rate of neurocognitive impairment (32) and lower risk of progression to AIDS (33) in PWH. CB consists of a heterogeneous mixture of compounds, but the primary cannabinoid constituent is Δ-9-tetrahydrocannabinol (THC). THC has been shown to suppresses expression of pro-inflammatory cytokines and thereby reduces systemic immune activation in HIV infection (34), but THC’s impact on the latent HIV reservoir is unclear.

However, recently published reports have indicated that reservoir size could be affected by CB use. Our lab recently showed, using a cohort of CB using PWH on ART, the CB use is associated with altered abundances of T cell subsets as well as a trend towards a smaller intact HIV reservoir (14). Another study also recently reported that CB use was associated with lower tissue reservoir burden in PWH with clade C infections (3).

In this present study, we have analyzed the impact of THC on HIV infection using a primary CD4 T cell model of HIV latency. Overall, we find that HIV expression and reactivation by LRAs are largely unaffected by THC exposure over a time period of 24 hours. Similarly, we saw no significant impact of THC on expression of the activation marker CD38 or on the expression of surface markers use to describe CD4 T cell memory subsets. This observation was robust across a wide range of THC concentrations. Nevertheless, it remains possible that a longer term or repeated exposure could influence HIV expression or LRA sensitivity. THC sensitivity of infected cells could also be modulated by the more complex environment *in vivo*, either in peripheral blood or in solid tissues such as the gut or brain. Animal models of HIV infection and CB exposure may thus be necessary to fully evaluate the impact of THC on HIV expression and reactivation.

In addition to measuring the impact of THC on HIV expression, we carried out an integrated single cell multiomic analysis of genome-wide chromatin accessibility and gene expression to characterize latently infected CD4 T cells exposed to THC. While these data showed only a small reduction in viral RNA and proviral accessibility after 6h of THC exposure, we observed a number of changes to gene expression and TF activity in HIV infected CD4 T cells. In particular, we observed a strong transcriptional downregulation of genes that contribute to protein synthesis, as well as interferon-stimulated genes. This observation suggests that THC represses macromolecular metabolism and protein expression in infected cells, which could affect the ability of infected cells to respond to antigen or cytokine cues. This finding thus could help to explain why CB users may have a smaller HIV reservoir size than non-users – if periodic T cell activation is necessary for intermittent clonal expansion that sustains the reservoir, THC suppression of T cell activation and metabolism could lead to a lower rate of reservoir clonal expansion over time, leading to more rapid reservoir depletion. The reduced expression of ISGs in infected cells also merits further investigation. The reduced activity of an innate sensing pathway in infected cells could contribute to the overall lower level of immune activation and exhaustion in PWH that use CB. The precise innate pathway that is active in infected cells and the molecular mechanism by which THC blocks this pathway will need to be elucidated.

Our analysis of differentially active transcription factors in HIV infected CD4 T cells after THC stimulation also produced some intriguing observations. Specifically, we observed that THC-induced increased binding activity of the transcriptional regulator CTCF. CTCF is a chromatin-binding protein that mediates long-range chromatin looping as part of the cohesin complex (35). CTCF has also been shown to silence the HIV promoter by inducing repressive chromatin structures (36). We also observed a strong enrichment of ETS TFs within the set of TFs with downregulated activity after THC exposure. Notably, ETS1 has been identified as a master regulator of ribosomal gene expression (37), suggesting that reduced activity of ETS transcription factors likely contribute to the reduced expression of ribosomal genes after THC exposure. We have also recently identified ETS1 as a key mediator of HIV repression in resting CD4 T cells (38). ETS family of transcription factors can act as activators or repressors of gene expression (38). THC exposure affects MAPK signaling pathways (39), which can regulate ETS family activity. For instance, the ERK pathway influenced by THC can modulate ETS1 or ELK3 activity, potentially affecting HIV latency dynamics. In addition to CTCF and ETS family of TF motifs, we also identified downregulated activity of Interferon regulator factor (IRF) proteins (IRF3, IRF4, IRF8, IRF9) in THC exposed HIV infected cells. Notably, this downregulation was not observed in uninfected CD4 T cells, suggesting that this phenomenon is specific to HIV infected cells. IRFs play key roles in the activation of ISGs in response to viral infection, consistent with our observation of reduced ISG expression in THC exposed cells. Further work to clarify the role of specific ETS transcription factors and IRFs in the response to THC in infected cells will be required.

Our findings should be considered in the light of several caveats and limitations to our study. Our approach uses an *in vitro* model of HIV latency that may not fully reflect the regulation of the clinical reservoir. Furthermore, CB is comprised of several cannabinoids other than THC that may play important roles in affecting the latent HIV reservoir and in immune exhaustion in PWH on ART. A broader investigation of the impact of minor cannabinoids on HIV infection could yield important new insights. The transcriptional and epigenomic response of infected cells to THC exposure could also vary depending on the length of exposure.

Nevertheless, our findings reveal some important new details regarding how CB use could impact the HIV reservoir and immune systems of PWH and highlight the value of multiomic approaches to understanding the responses of immune cells to stimuli. In particular, the downregulation of protein synthesis and antiviral pathways in HIV infected cells following THC exposure could be a significant contributor to the documented impact of CB use on the immune systems of PWH. Further research will help to clarify these connections and potentially help to guide HIV cure approaches that are customized for PWH with substance abuse disorder.

## Author contributions

**Manickam Ashokkumar:** Methodology, Validation, Formal analysis, Investigation, Data curation, Writing – original draft, Visualization. **Renee Y Ge:** Methodology, Software, Formal analysis, Data curation, Writing – review & editing, Visualization. **Alicia Cooper-Volkheimer:** Methodology, Investigation, Data curation, Writing – review & editing. **David M. Margolis:** Methodology, Resources, Funding acquisition, Writing – review & editing. **Quefeng Li:** Methodology, Software, Formal analysis, Resources, Data curation, Visualization, Supervision, Writing – review & editing. **Yuchao Jiang:** Methodology, Software, Formal analysis, Resources, Data curation, Visualization, Supervision, Project administration. Funding acquisition, Writing – review & editing. **David M. Murdoch:** Methodology, Resources, Supervision. Funding acquisition, Writing – review & editing. **Edward P. Browne:** Conceptualization, Methodology, Validation, Formal analysis, Resources, Writing – original draft, Supervision, Project administration, Funding acquisition. All authors have read and approved the final manuscript.

## Acknowledgement

We thank the UNC HIV cure center, Center for AIDS Research, the UNC Flow Cytometry Core Facility, and the Fred Hutch Bioinformatics and Genomics Cores. This work was supported by the following grants from the National Institutes of Health: R35 GM138342 (YJ), R61 DA053599 (EPB), R61 DA059918 (EPB), and UM1 AI164567 (DMM).

---

**Table S1. Differentially expressed genes in immune cells after THC exposure.**

**Table S2. Enrichment analysis of downregulated genes from the uninfected CD4 T cell clusters using MSigDB Hallmark gene sets.**

**Table S3. Enrichment analysis of downregulated genes from the infected CD4 T cell clusters using MSigDB Hallmark gene sets.**

**Table S4. Gene set enrichment analysis of downregulated genes from the uninfected CD4 T cell clusters using Gene ontology.**

**Table S5. Gene set enrichment analysis of downregulated genes from the infected CD4 T cell clusters using Gene ontology.**

**Table S6. Differentially accessible transcription factors in immune cells after THC exposure.**

**Supplementary Figure S1. Impact of THC on cell surface marker phenotype and HIV expression of latently infected primary CD4 cells at 6h.**

Latently infected CD4 cells were exposed to THC at a range of concentrations for 6 h. At 6 h post exposure, we quantified the distribution of different CD4 T cell subsets, expression of activation marker, CD38, and HIV (GFP) expression. (**A**) Schematic of latently infected CD4+ T cell generation, pooling and THC stimulation followed by flowcytometry. (**B**) Abundance of CD4 T cell subsets, naïve (Tn), central memory (Tcm), effector memory (Tem) and effector (Teff) subsets at 6 h post THC administration/exposure. (**C**) Frequency of CD38+ cells within CD4 T cells subsets (Tn, Tcm, Tem, and Teff cells) after THC exposure. (**D**) Expression of GFP (HIV) within CD4 T cells subsets (Tn, Tcm, Tem, and Teff cells) at 6h of THC exposure. Cell subsets were assigned by expression of the phenotype surface markers, CD45RA and CCR7 – Tn: CD45RA+/CCR7+, Tcm: CD45RA-/CCR7+, Tem: CD45RA-/CCR7-, Teff: CD45RA+/CCR7-. None of the sample achieved statistical significance compared to control treated cells.

**Supplementary Figure S2. UMAP dimension reduction plots with colors corresponding to HIV vRNA expression and ATAC accessibility.**

**(A)** Violin plot of the scRNA-seq data based on the level of HIV vRNA expression without (left panel) and with (right panel) THC exposure for 6h. **(B)** Violin plot of the scATAC-seq data based on the level of HIV ATAC accessibility without (left panel) and with (right panel) THC exposure for 6 h.

## References

1. Cohn LB, Chomont N, Deeks SG. The Biology of the HIV-1 Latent Reservoir and Implications for Cure Strategies. Cell Host Microbe. 2020;27(4):519–30.

2. Manuzak JA, Gott TM, Kirkwood JS, Coronado E, Hensley-McBain T, Miller C, et al. Heavy Cannabis Use Associated With Reduction in Activated and Inflammatory Immune Cell Frequencies in Antiretroviral Therapy-Treated Human Immunodeficiency Virus-Infected Individuals. Clin Infect Dis. 2018;66(12):1872–82.

3. Liu Z, Julius P, Himwaze CM, Mucheleng’anga LA, Chapple AG, West JT, et al. Cannabis Use Associates With Reduced Proviral Burden and Inflammatory Cytokine in Tissues From Men With Clade C HIV-1 on Suppressive Antiretroviral Therapy. The Journal of Infectious Diseases. 2024;229(5):1306–16.

4. Min AK, Keane AM, Weinstein MP, Swartz TH. The impact of cannabinoids on inflammasome signaling in HIV-1 infection. NeuroImmune Pharm Ther. 2023;2(1):79–88.

5. Kumar V, Torben W, Mansfield J, Alvarez X, Vande Stouwe C, Li J, et al. Cannabinoid Attenuation of Intestinal Inflammation in Chronic SIV-Infected Rhesus Macaques Involves T Cell Modulation and Differential Expression of Micro-RNAs and Pro-inflammatory Genes. Front Immunol. 2019;10:914.

6. Roth MD, Tashkin DP, Whittaker KM, Choi R, Baldwin GC. Tetrahydrocannabinol suppresses immune function and enhances HIV replication in the huPBL-SCID mouse. Life Sci. 2005;77(14):1711–22.

7. Molina PE, Amedee A, LeCapitaine NJ, Zabaleta J, Mohan M, Winsauer P, et al. Cannabinoid neuroimmune modulation of SIV disease. J Neuroimmune Pharmacol. 2011;6(4):516–27.

8. Nagarkatti P, Pandey R, Rieder SA, Hegde VL, Nagarkatti M. Cannabinoids as novel anti-inflammatory drugs. Future Med Chem. 2009;1(7):1333–49.

9. Cabral GA, Griffin-Thomas L. Emerging role of the cannabinoid receptor CB2 in immune regulation: therapeutic prospects for neuroinflammation. Expert Rev Mol Med. 2009;11:e3.

10. McCoy KL. Interaction between Cannabinoid System and Toll-Like Receptors Controls Inflammation. Mediators of Inflammation. 2016;2016(1):5831315.

11. Howlett AC, Blume LC, Dalton GD. CB(1) cannabinoid receptors and their associated proteins. Curr Med Chem. 2010;17(14):1382–93.

12. Zou S, Kumar U. Cannabinoid Receptors and the Endocannabinoid System: Signaling and Function in the Central Nervous System. International Journal of Molecular Sciences. 2018;19(3):833.

13. Samson M-T, Small-Howard A, Shimoda LMN, Koblan-Huberson M, Stokes AJ, Turner H. Differential Roles of CB1 and CB2 Cannabinoid Receptors in Mast Cells1. The Journal of Immunology. 2003;170(10):4953–62.

14. Falcinelli SD, Cooper-Volkheimer AD, Semenova L, Wu E, Richardson A, Ashokkumar M, et al. Impact of Cannabis Use on Immune Cell Populations and the Viral Reservoir in People With HIV on Suppressive Antiretroviral Therapy. J Infect Dis. 2023;228(11):1600–9.

15. Klein TW, Newton C, Larsen K, Lu L, Perkins I, Nong L, et al. The cannabinoid system and immune modulation. J Leukoc Biol. 2003;74(4):486–96.

16. Ashokkumar M, Hafer TL, Felton A, Archin NM, Margolis DM, Emerman M, et al. A targeted CRISPR screen identifies ETS1 as a regulator of HIV latency. bioRxiv. 2024:2024.08.03.606477.

17. Ashokkumar M, Mei W, Peterson JJ, Harigaya Y, Murdoch DM, Margolis DM, et al. Integrated Single-cell Multiomic Analysis of HIV Latency Reversal Reveals Novel Regulators of Viral Reactivation. Genomics, Proteomics & Bioinformatics. 2024.

18. Germain PL, Lun A, Garcia Meixide C, Macnair W, Robinson MD. Doublet identification in single-cell sequencing data using scDblFinder. F1000Res. 2021;10:979.

19. Stuart T, Butler A, Hoffman P, Hafemeister C, Papalexi E, Mauck WM, 3rd, et al. Comprehensive Integration of Single-Cell Data. Cell. 2019;177(7):1888–902.e21.

20. Stuart T, Srivastava A, Madad S, Lareau CA, Satija R. Single-cell chromatin state analysis with Signac. Nat Methods. 2021;18(11):1333–41.

21. Jiang Y, Harigaya Y, Zhang Z, Zhang H, Zang C, Zhang NR. Nonparametric single-cell multiomic characterization of trio relationships between transcription factors, target genes, and cis-regulatory regions. Cell Syst. 2022;13(9):737–51.e4.

22. Becht E, McInnes L, Healy J, Dutertre C-A, Kwok IWH, Ng LG, et al. Dimensionality reduction for visualizing single-cell data using UMAP. Nature Biotechnology. 2019;37(1):38–44.

23. Guan PY, Lee JS, Wang L, Lin KZ, Mei W, Chen L, et al. Destin2: Integrative and cross-modality analysis of single-cell chromatin accessibility data. Frontiers in Genetics. 2023;Volume 14 - 2023.

24. Fornes O, Castro-Mondragon JA, Khan A, van der Lee R, Zhang X, Richmond PA, et al. JASPAR 2020: update of the open-access database of transcription factor binding profiles. Nucleic Acids Res. 2020;48(D1):D87–d92.

25. Schep AN, Wu B, Buenrostro JD, Greenleaf WJ. chromVAR: inferring transcription-factor-associated accessibility from single-cell epigenomic data. Nature Methods. 2017;14(10):975–8.

26. Kuleshov MV, Jones MR, Rouillard AD, Fernandez NF, Duan Q, Wang Z, et al. Enrichr: a comprehensive gene set enrichment analysis web server 2016 update. Nucleic Acids Res. 2016;44(W1):W90–7.

27. Camacho-Pereira J, Tarragó MG, Chini CCS, Nin V, Escande C, Warner GM, et al. CD38 Dictates Age-Related NAD Decline and Mitochondrial Dysfunction through an SIRT3-Dependent Mechanism. Cell Metab. 2016;23(6):1127–39.

28. Tsompana M, Buck MJ. Chromatin accessibility: a window into the genome. Epigenetics & Chromatin. 2014;7(1):33.

29. Suryavanshi SV, Kovalchuk I, Kovalchuk O. Cannabinoids as Key Regulators of Inflammasome Signaling: A Current Perspective. Front Immunol. 2020;11:613613.

30. Maggirwar SB, Khalsa JH. The Link between Cannabis Use, Immune System, and Viral Infections. Viruses. 2021;13(6).

31. Yin L, Dinasarapu AR, Borkar SA, Chang KF, De Paris K, Kim-Chang JJ, et al. Anti-inflammatory effects of recreational marijuana in virally suppressed youth with HIV-1 are reversed by use of tobacco products in combination with marijuana. Retrovirology. 2022;19(1):10.

32. Watson CW, Paolillo EW, Morgan EE, Umlauf A, Sundermann EE, Ellis RJ, et al. Cannabis Exposure is Associated With a Lower Likelihood of Neurocognitive Impairment in People Living With HIV. J Acquir Immune Defic Syndr. 2020;83(1):56–64.

33. Coates RA, Farewell VT, Raboud J, Read SE, MacFadden DK, Calzavara LM, et al. Cofactors of progression to acquired immunodeficiency syndrome in a cohort of male sexual contacts of men with human immunodeficiency virus disease. Am J Epidemiol. 1990;132(4):717–22.

34. Mboumba Bouassa RS, Comeau E, Alexandrova Y, Pagliuzza A, Yero A, Samarani S, et al. Effects of Oral Cannabinoids on Systemic Inflammation and Viral Reservoir Markers in People with HIV on Antiretroviral Therapy: Results of the CTN PT028 Pilot Clinical Trial. Cells. 2023;12(14).

35. Zhou R, Tian K, Huang J, Duan W, Fu H, Feng Y, et al. CTCF DNA-binding domain undergoes dynamic and selective protein-protein interactions. iScience. 2022;25(9):105011.

36. Jefferys SR, Burgos SD, Peterson JJ, Selitsky SR, Turner A-MW, James LI, et al. Epigenomic characterization of latent HIV infection identifies latency regulating transcription factors. PLOS Pathogens. 2021;17(2):e1009346.

37. Xiao FH, Yu Q, Deng ZL, Yang K, Ye Y, Ge MX, et al. ETS1 acts as a regulator of human healthy aging via decreasing ribosomal activity. Sci Adv. 2022;8(17):eabf2017.

38. Ashokkumar M, Hafer TL, Felton A, Archin NM, Margolis DM, Emerman M, et al. A targeted CRISPR screen identifies ETS1 as a regulator of HIV-1 latency. PLoS Pathog. 2025;21(4):e1012467.

39. Simon L, Song K, Vande Stouwe C, Hollenbach A, Amedee A, Mohan M, et al. Δ9-Tetrahydrocannabinol (Δ9-THC) Promotes Neuroimmune-Modulatory MicroRNA Profile in Striatum of Simian Immunodeficiency Virus (SIV)-Infected Macaques. J Neuroimmune Pharmacol. 2016;11(1):192–213.

